# Major role of IgM in the neutralizing activity of convalescent plasma against SARS-CoV-2

**DOI:** 10.1101/2020.10.09.333278

**Authors:** Romain Gasser, Marc Cloutier, Jérémie Prévost, Corby Fink, Éric Ducas, Shilei Ding, Nathalie Dussault, Patricia Landry, Tony Tremblay, Audrey Laforce-Lavoie, Antoine Lewin, Guillaume Beaudoin-Bussières, Annemarie Laumaea, Halima Medjahed, Catherine Larochelle, Jonathan Richard, Gregory A. Dekaban, Jimmy D. Dikeakos, Renée Bazin, Andrés Finzi

## Abstract

Characterization of the humoral response to SARS-CoV-2, the etiological agent of Covid-19, is essential to help control the infection. In this regard, we and others recently reported that the neutralization activity of plasma from COVID-19 patients decreases rapidly during the first weeks after recovery. However, the specific role of each immunoglobulin isotype in the overall neutralizing capacity is still not well understood. In this study, we selected plasma from a cohort of Covid-19 convalescent patients and selectively depleted immunoglobulin A, M or G before testing the remaining neutralizing capacity of the depleted plasma. We found that depletion of immunoglobulin M was associated with the most substantial loss of virus neutralization, followed by immunoglobulin G. This observation may help design efficient antibody-based COVID-19 therapies and may also explain the increased susceptibility to SARS-CoV-2 of autoimmune patients receiving therapies that impair the production of IgM.

## Introduction

Since its discovery in Wuhan in 2019, the causative agent of COVID-19, the SARS-CoV-2 virus (Zhu et al., 2020), has become a major global public health problem. A better understanding of immune responses induced by SARS-CoV-2 is urgently needed to help control the infection. Several studies have shown that the neutralization activity of plasma from COVID-19 patients decreases rapidly during the first weeks after recovery (Beaudoin-Bussières et al., 2020; Long et al., 2020; Prévost et al., 2020; Robbiani et al., 2020; Seow et al., 2020). Although a good correlation between the presence of Spike (S)-specific antibodies and the capacity of plasma from infected individuals to neutralize viral particles was reported, recent data looking at individual immunoglobulin (Ig) isotypes revealed a stronger correlation between the decrease in S-specific IgM antibodies and loss of neutralization compared to S-specific IgG and IgA antibodies, suggesting that IgM play an important role in the neutralization activity of plasma from individuals who suffered from COVID-19 (Beaudoin-Bussières et al., 2020; Prévost et al., 2020). To better understand the relative contribution of S-specific IgM, IgA and IgG antibodies in SARS-CoV-2 neutralization, we selectively depleted each Ig isotype from plasma obtained from 25 convalescent donors and assessed the impact of depletion on the capacity of the plasma to neutralize SARS-CoV-2 pseudoviral particles and wild type infectious SARS-CoV-2 viral particles.

## Results

### Ig depletion

Demographic information of the 25 convalescent donors (21 males, 4 females, median = 45 days after symptoms onset), who were diagnosed with or tested positive for SARS-CoV-2 with complete resolution of symptoms for at least 14 days before sampling are presented in Table 1. Selective depletion of IgM, IgA or IgG was achieved by adsorption on isotype-specific ligands immobilized on Sepharose or agarose beads, starting with a five-fold dilution of plasma (see details in Stars Methods). The depletion protocols permitted to efficiently deplete each isotype while leaving the other isotypes nearly untouched, as measured by ELISA (Fig 1A-C). Depletion of IgG had a much higher impact on the total level of SARS-CoV-2 RBD antibodies than IgM and IgA depletion (Fig 1D), although RBD-specific antibodies of each isotype were selectively removed by the depletion (Fig. 1E-G). The impact of IgG depletion on the level of total antibodies against the full S glycoprotein expressed on 293T cells (measured by flow cytometry) was also noticeable (Fig. 1H) whereas isotype-specific detection of full S antibodies by flow cytometry confirmed the efficacy of selective depletion (Fig. 1I-K).

**Table 1.** COVID convalescent plasma donor’s characteristics.

|  | All donors | Males | Females |
| --- | --- | --- | --- |
| Donors (n) | 25 | 21 | 4 |
| Average age ± SD [range] | 47 ± 16 [20-69] | 49 ± 17 [20-69] | 40 ± 14 [29-60] |
| Age (median) | 50 | 51 | 34.5 |
| Period (days) between symptoms onset and donation (median [range]) | 45 [25-69] | 47 [25-69] | 40 [27-56] |

**Figure 1.**
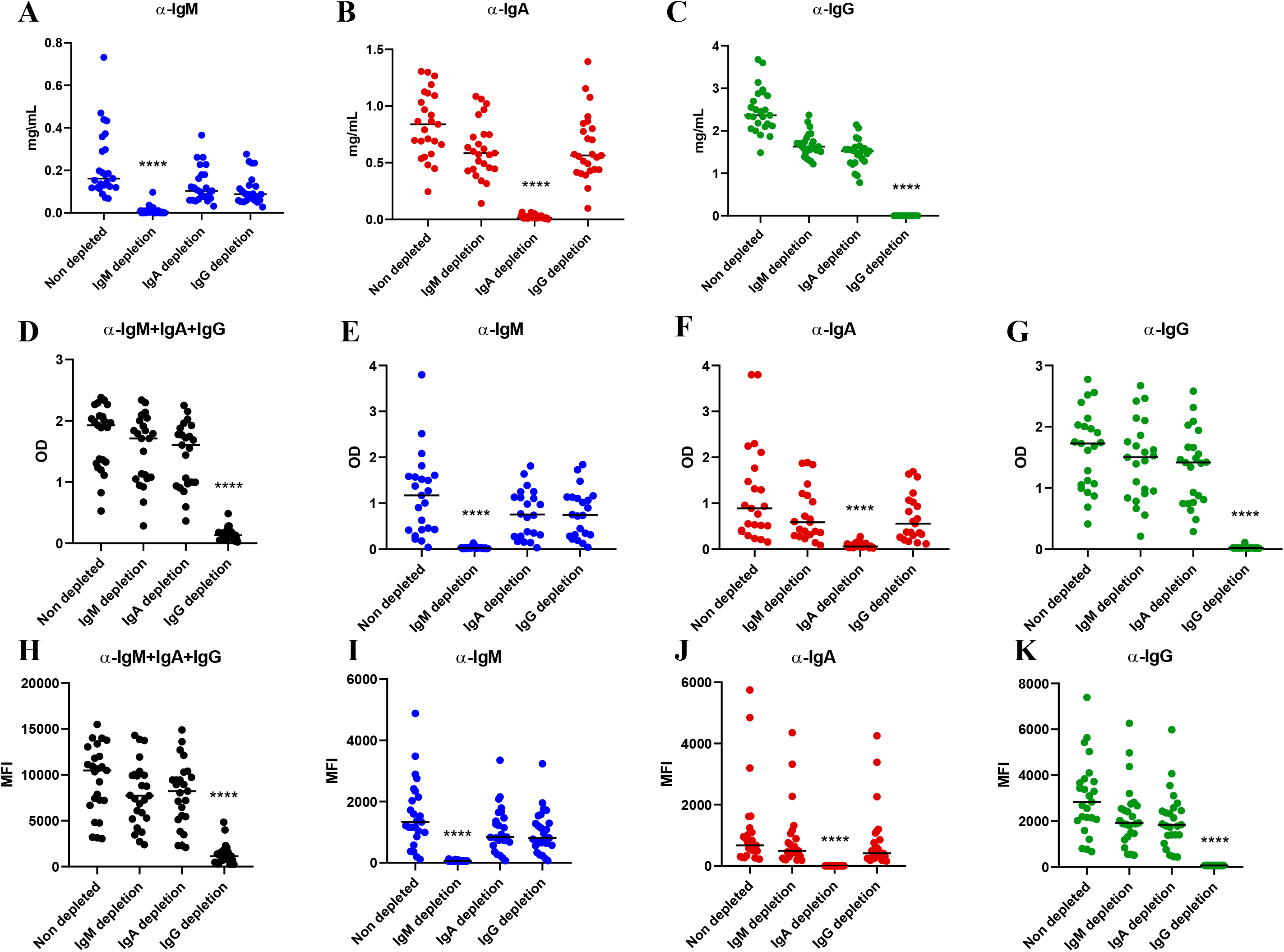
IgM, IgA and IgG depletion in plasma samples from convalescent donors. **(A-C)** Efficacy of the specific isotype depletion assessed by ELISA for total IgM, IgA and IgG. All plasma samples were diluted 5-fold prior to depletion; (A) IgM concentration in non-depleted, IgM-depleted, IgA-depleted and IgG-depleted plasmas, measured using an anti-human IgM (μ-chain specific) as capture antibody; (B) IgA concentration measured on the same plasmas using anti-human IgA (α-chain specific); (C) IgG concentration measured using anti-human IgG (γ-chain specific). (**D-G)** Efficacy of SARS-CoV-2 specific antibody depletion assessed by SARS-CoV-2 RBD ELISA; (D) Level of total (pan-Ig) anti-SARS-CoV-2 RBD-specific antibodies in non-depleted, IgM-depleted, IgA depleted and IgG-depleted plasmas; (E) Level of IgM-specific anti-RBD; (F) Level of IgA-specific anti-RBD; (G) Level of IgG-specific anti-RBD. (**H-K)** Efficacy of full S glycoprotein-specific antibody depletion measured by flow cytometry; (H) Level of total (pan-Ig) anti-SARS-CoV-2 S-specific antibodies in non-depleted, IgM-depleted, IgA-depleted and IgG-depleted plasmas; (I) Level of IgM-specific anti-S; (J) Level of IgA-specific anti-S; (K) Level of IgG-specific anti-S. Asterisks indicate the level of statistical significance obtained by a Dunn’s test; **** p<0.0001.

### Neutralizing activity of depleted plasma

We then evaluated the capacity of non-depleted and isotype-depleted plasma samples to neutralize pseudoviral particles expressing the S glycoprotein from SARS-CoV-2 (Prévost et al., 2020) (Star Methods). Depletion of IgM, IgA or IgG all resulted in a significant decrease of neutralization compared to non-depleted plasma (Fig. 2A-D). However, the loss of neutralization activity was much more pronounced in IgM- and IgG-depleted plasma with a 5.5 and 4.5 fold decrease in mean ID_50_ compared to non-depleted plasma respectively, than in IgA-depleted plasma where a 2.4 fold decrease only was observed (Fig. 2E). To evaluate whether the impact of isotype depletion on neutralization could be extended beyond pseudoviral particles, we tested plasma from eight donors in microneutralization experiments using fully infectious SARS-CoV-2 viral particles, as described in the Star Methods. The neutralizing potency of plasma was greatly reduced following IgM and IgG (4.0 and 2.9 fold respectively) but not IgA (no decrease) depletion (Fig. 2F and G). Despite the limited number of samples tested with the live virus, the impact of IgM and IgG depletion on neutralization was similar to that observed with the same samples in the pseudoviral particles neutralization assay (Fig. 3A-C). This data not only confirms the role of IgG in neutralizing activity of convalescent plasma but also highlights the important contribution of IgM with respect to neutralization activity.

**Figure 2.**
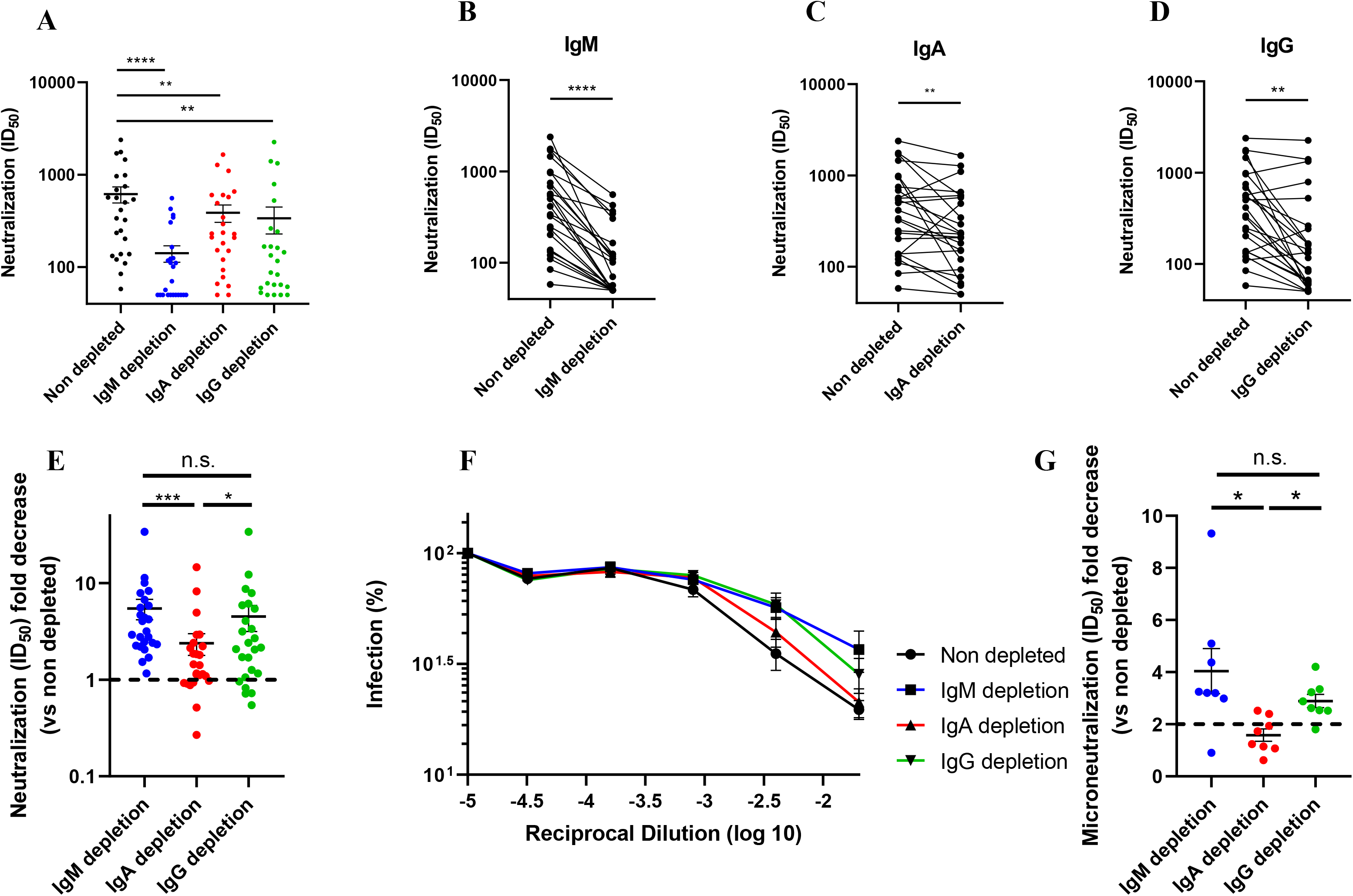
Role of IgM, IgA and IgG in neutralization. **(A)** Comparison of the SARS-CoV-2 pseudoviral inhibitory dilution (ID_50_) of all plasma samples. **(B-D)** ID_50_ of plasma from each convalescent donor before and after (B) IgM, (C) IgA and (D) IgG depletion. **(E)** Fold decrease (isotype-depleted versus non-depleted plasma) in ID_50_ measured by SARS-CoV-2 pseudovirions neutralization. **(F-G)** Microneutralization assay using infectious wild type SARS-CoV-2 performed on non-depleted and isotype-depleted plasma from 3 donors; (F) Mean percentage of infection observed with plasma from the 3 donors and (G) Fold decrease (isotype-depleted versus non-depleted plasma) in ID_50_ measured by microneutralization of wild type SARS-CoV-2 virions. Asterisks indicate the level of statistical significance obtained by a Wilcoxon signed rank test, n.s. not significant; *p<0.05; **p<0.01; ***p<0.001; ****p<0.0001.

**Figure 3.**
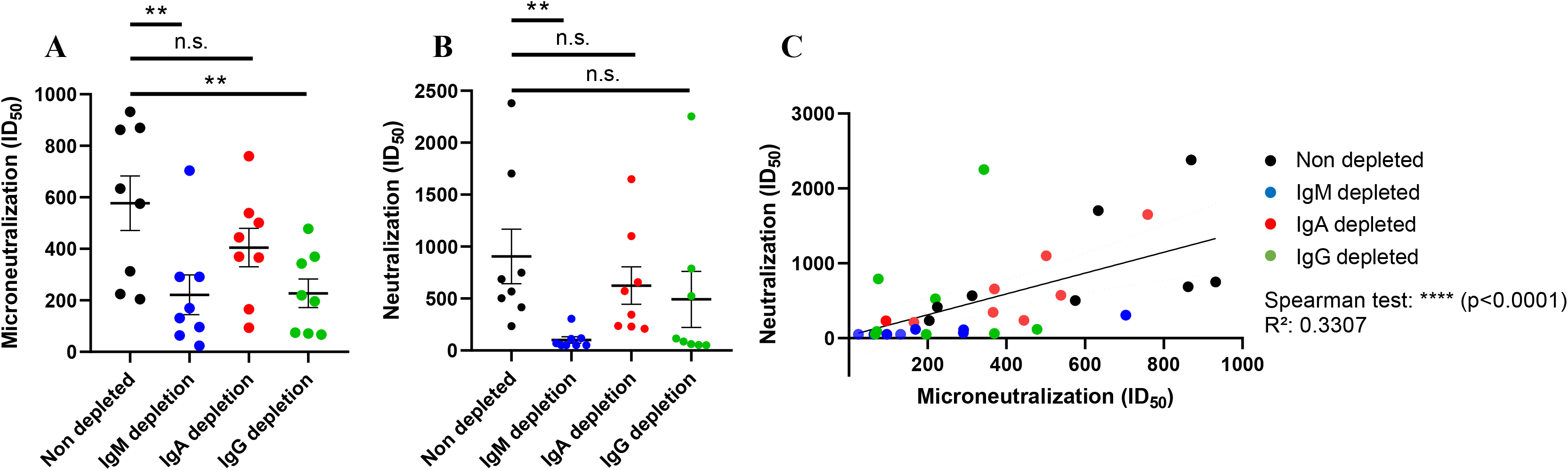
Neutralizing capacity of eight convalescent plasma using pseudoviral particles or microneutralization with infectious wild type SARS-CoV-2 virus. ID_50_ obtained by (**A**) virus microneutralization assay or (**B**) pseudoviral particles neutralization assay for non-depleted or isotype-depleted plasma of eight convalescent donors. (**C**) Spearman correlation and linear regression fitting between the ID_50_ obtained by microneutralization and pseudoviral particles neutralization assays. Dashed lines indicate the 95% confidence interval of the linear regression fitting. Non-depleted plasmas are shown in black, IgM-depleted in blue, IgA-depleted in red and IgG-depleted in green. Asterisks indicate the level of statistical significance obtained by a Wilcoxon signed rank test, n.s. not significant; **p<0.01; ****p<0.0001.

## Discussion

Our findings detailing the important role of IgM in the neutralizing activity of convalescent plasma has several implications. First, although the therapeutic efficacy of convalescent plasma for the treatment of COVID-19 patients remains to be established, it is likely that neutralizing antibodies will play a role. Because SARS-CoV-2 specific IgM antibodies rapidly decrease after disease onset (Beaudoin-Bussières et al., 2020; Prévost et al., 2020; Robbiani et al., 2020; Seow et al., 2020), the collection of convalescent plasma with maximal neutralizing activity should be performed early after disease recovery. Second, our results suggest that caution should be taken when using therapeutics that impair the production of IgM. Anti-CD20 antibodies (B cell-depleting agents) are used to treat several inflammatory disorders. Their use is associated with IgM deficiency in a substantial number of patients, while their impact on IgG and IgA levels is more limited (Kridin and Ahmed, 2020). In line with our data, recent studies reported that anti-CD20 therapy could be associated with a higher susceptibility to contract SARS-CoV-2 and develop severe COVID-19 (Guilpain et al., 2020; Hughes et al., 2020; Safavi et al., 2020; Schulze-Koops et al., 2020; Sharmeen et al., 2020; Sormani et al., 2020). Whether this is associated to the preferential depletion of IgM-producing B cells by these treatments (Looney et al., 2008) remains to be shown. Nevertheless, our results suggest that IgM levels should be investigated as a biomarker to stratify patients on immunosuppressive therapies at higher risk for COVID-19.

In summary, our results extend previous observations showing a strong correlation between neutralization potency and the presence of RBD-specific IgM (Beaudoin-Bussières et al., 2020; Perera et al., 2020; Prévost et al., 2020; Seow et al., 2020). It is intriguing that IgM represents about only 5% of the total antibodies in plasma (Wang et al., 2020), yet plays such an important role in SARS-CoV-2 neutralization. Whether this is due to the enhanced avidity provided by its pentameric nature remains to be formally demonstrated but is in agreement with recent work demonstrating that dimeric antibodies are more potent than their monomeric counterpart (Wang et al., 2020). The possible establishment of long lived IgM-producing B cells that might contribute to long term immunity of recovered patients has been suggested (Brouwer et al., 2020; Newell et al., 2020). However, how plasma neutralization evolves over prolonged periods of time and the specific role of IgM in this activity remains to be determined.

## Acknowledgments

This work was supported by “Ministère de l’Économie et de l’Innovation du Québec, Programme de soutien aux organismes de recherche et d’innovation”, by the Fondation du CHUM, by the Canada’s COVID-19 Immunity Task Force (CITF), in collaboration with the Canadian Institutes of Health Research (CIHR) and a CIHR foundation grant #352417 to A.F. Funding was also provided by an operating grant from CIHR from the Canadian 2019 Novel Coronavirus (COVID-19) Rapid Research Funding Opportunity (FRN440388 to JDD and GAD) and an Infrastructure Grant from CFI for the Imaging Pathogens for Knowledge Translation (ImPaKT) Facility (#36287 to JDD and GAD). A.F. is the recipient of a Canada Research Chair on Retroviral Entry # RCHS0235 950-232424. R.G. is supported by a MITACS Accélération postdoctoral fellowship. J.P. is supported by a CIHR graduate fellowship.

## Author Contributions

R.G., M.C., J.P., R.B. and A.F. designed the studies. R.G. and S.D. performed neutralization experiments with pseudoviral particles. J.P. performed flow cytometry experiments. C.F., G.A.D. and J.D.D. performed microneutralization assays with infectious wildtype SARS-CoV-2 and analysed the results. M.C., E.D., N.D., P.L., A.L.L. and T.T. depleted plasma samples and performed the ELISA. J.R. provided new reagents. A.L. performed statistical analysis. C.L. provided scientific and clinical input. R.G., M.C., R.B. and A.F. wrote the manuscript with inputs from others. Every author has read, edited and approved the final manuscript.

## Material and Methods

### Ethics statement

All work was conducted in accordance with the Declaration of Helsinki in terms of informed consent and approval by an appropriate Ethics Review board. Convalescent plasmas were obtained from donors who consented to participate in this research project at CHUM (19.381) and at Héma-Québec (REB # 2020-004). The donors met all donor eligibility criteria: previous confirmed COVID-19 infection and complete resolution of symptoms for at least 14 days.

### Plasmids

The plasmids expressing the human coronavirus Spike of SARS-CoV-2 was kindly provided by Stefan Pöhlmann and was previously reported (Hoffmann et al., 2020). The pNL4.3 R-E-Luc was obtained from NIH AIDS Reagent Program. The codon-optimized RBD sequence (encoding residues 319-541) fused to a C-terminal hexahistidine tag was cloned into the pcDNA3.1(+) expression vector and was reported elsewhere (Beaudoin-Bussières et al., 2020). The vesicular stomatitis virus G (VSV-G)-encoding plasmid (pSVCMV-IN-VSV-G) was previously described (Lodge et al., 1997).

### Cell lines

293T human embryonic kidney cells (obtained from ATCC) and Vero E6 cells (ATCC CRL-1586™) were maintained at 37°C under 5% CO2 in Dulbecco’s modified Eagle’s medium (DMEM) (Wisent) containing 5% fetal bovine serum (VWR), 100 UI/ml of penicillin and 100μg/ml of streptomycin (Wisent). The 293T-ACE2 cell line was previously reported (Prévost et al., 2020). For the generation of 293T cells stably expressing SARS-CoV-2 Spike, VSV-G pseudotyped lentivirus packaging the SARS-CoV-2 Spike was produced in 293T using a third-generation lentiviral vector system. Briefly, 293T cells were co-transfected with two packaging plasmids (pLP1 and pLP2), an envelope plasmid (pSVCMV-IN-VSV-G) and a lentiviral transfer plasmid coding for a GFP-tagged SARS-CoV-2 Spike (pLV-SARS-CoV-2 S C-GFPSpark tag) (SinoBiological). Supernatant containing lentiviral particles was used to infect 293T cells in presence of 5μg/mL polybrene. The 293T cells stably expressing SARS-CoV-2 Spike (GFP+) were sorted by flow cytometry. SARS-CoV-2 expression was confirmed using the CR3022 mAb and plasma from SARS-CoV-2-infected individuals.

### Isotype depletion

Selective depletion of IgM, IgA or IgG was done by adsorption on isotype-specific ligands immobilized on sepharose or agarose beads starting with a five-fold dilution of plasma in PBS. IgG and IgA antibodies were depleted from plasma obtained from 25 recovered COVID-19 patient using Protein G HP Spintrap (GE Healthcare Life Sciences, Buckinghamshire, UK) and Peptide M / Agarose (InvivoGen, San Diego, CA), respectively, according to the manufacturer’s instructions with the exception that no elution step for the recovery of the targeted antibodies was done. For IgM depletion, anti-human IgM (μ-chain specific, Sigma, St. Louis, MO) was covalently coupled to NHS HP SpinTrap (GE Healthcare) at 815 μg/mL of matrix. Depletion was performed according to the manufacturer’s instructions with the exception that no elution step for the recovery of the targeted isotype was done. All non-depleted and isotype-depleted samples were filtered on a 0.22 μm Millex GV filter (SLGV013SL, Millipore, Burlington, MA) to ensure sterility for the virus capture and neutralization assays.

### Immunoglobulin isotype ELISA

To assess the extent of IgM, IgG and IgA depletion, ELISA were performed on non-depleted as well as IgM-, IgA- and IgG-depleted plasma samples. Each well of a 96-well microplate was filled with either goat anti-human IgM (μ-chain specific) at 5 μg/mL, goat anti-human serum IgA (α-chain specific) at 0.3 μg/mL or goat anti-human IgG (γ-chain specific) at 5 μg/mL (all from Jackson ImmunoResearch Laboratories, Inc., West Grove, PA). Microtiter plates were sealed and stored overnight at 2-8°C. After four (IgA) to six (IgM and IgG) washes with H_2_O-0.1% Tween 20 (Sigma), 200 μL of blocking solution (10 mmol/L phosphate buffer, pH 7.4, containing 0.85% NaCl, 0.25% Hammerstein casein (EMD Chemicals Inc., Gibbstown, NJ,) were added to each well to block any remaining binding sites. The blocking solution for the IgG and IgM ELISA also contained 0.05% Tween 20. After 0.5 (IgA) to 1h (IgM and IgG) incubation at 37°C and washes, samples and the standard curves (prepared with human calibrated standard serum, Cedarlane, Burlington, Canada) were added to the plates in triplicates. Plates were incubated for 1h at 37°C. After washes, 100 μL of either goat anti-human IgA+G+M (H+L) HRP conjugate (1/30 000), goat anti-human IgG (H+L) HRP conjugate (1/30 000) or goat anti-human IgA (α-chain specific) HRP conjugate (1/10 000) (all from Jackson ImmunoResearch Laboratories, Inc.) were added and samples were incubated at 37°C for 1h. Wells were washed and bound antibodies were detected by the addition of 100 μL of 3,3′,5,5′-tetramethylbenzimidine (TMB, ScyTek Laboratories, Logan, UT). The enzymatic reaction was stopped by the addition of 100 μL 1 N H_2_SO_4_ and the absorbance was measured at 450/630 nm within 5 minutes.

### SARS-CoV-2 RBD ELISA

The presence of SARS-CoV-2 RBD-specific antibodies in the plasma from 25 recovered COVID-19 patients before and after depletion was measured using an ELISA adapted from a recently described protocol (Beaudoin-Bussières et al., 2020; Perreault et al., 2020; Prévost et al., 2020). The plasmid encoding for SARS-CoV-2 RBD was synthesized commercially (Genscript, Piscataway, NJ, USA). Recombinant RBD proteins were produced in transfected FreeStyle 293F cells (Invitrogen, Carlsbad, CA, USA) and purified by nickel affinity chromatography. Recombinant RBD was diluted to 2.5 μg/mL in PBS (Thermo Fisher Scientific, Waltham, MA, USA) and 100 μl of the dilution was distributed in the wells of flat-bottom 96-well microplates (Immulon 2HB; Thermo Scientific). The plates were placed overnight at 2-8°C for antigen adsorption. For the assay, the plates were emptied and a volume of 300 μl/well of blocking buffer (PBS-0.1% Tween (Sigma)-2% BSA (Sigma)) was added. The microplates were incubated for one hour at room temperature (RT) followed by washing four times (ELx405 microplate washer, Bio-Tek) with 300 μL/well of washing solution (PBS-0.1% Tween). Because the reaction is time sensitive, samples, negative and positive controls were prepared in triplicates in a plate, then transferred in the RBD coated plate by reverse multi-pipetting. The negative control was prepared from a pool of 23 COVID negative plasmas while the positive control was a characterized plasma from a recovered patient. After transfer, the plates were incubated for 60 minutes at 20-24°C. After four washes, 100 μL of either goat anti-human IgA+G+M (H+L) HRP conjugate (1/30 000) for the detection of all isotypes, goat anti-human IgM (μ-chain specific) HRP conjugate (1/15 000), F(ab’)₂ fragment goat anti-human IgA (α-chain specific) HRP conjugate (1/4500) (all from Jackson Immunosearch Laboratories, Inc.) or goat anti-human IgG (γ-chain specific) HRP conjugate (1/50 000) (Invitrogen) were added and samples were incubated at 20-24°C for 60 minutes. Wells were washed four times and bound antibodies were detected by the addition of 100 μL of 3,3′,5,5′-tetramethylbenzimidine (ScyTek Laboratories). The enzymatic reaction was stopped by the addition of 100 μL 1 N H_2_SO_4_ and the absorbance was measured at 450/630 nm within 5 minutes.

### Flow cytometry analysis of cell-surface staining

293T cells stably expressing SARS-CoV-2 Spike with a C-GFP tag (293T-Spike) were mixed at a 1:1 ratio with non-transduced 293T cells and were stained with plasma from SARS-CoV-2-infected individuals (1:250 dilution). Plasma binding to cell-surface Spike was revealed using fluorescent secondary antibodies able to detect all Ig isotypes (anti-human IgM+IgG+IgA; Jackson ImmunoResearch Laboratories, Inc.) or specific to IgG isotype (Biolegend), IgM isotype (Jackson ImmunoResearch Laboratories, Inc.) or IgA isotype (Jackson ImmunoResearch Laboratories, Inc.). The living cell population was gated on the basis of a viability dye staining (Aqua Vivid, Invitrogen). Samples were acquired on a LSRII cytometer (BD Biosciences, Mississauga, ON, Canada) and data analysis was performed using FlowJo v10.5.3 (Tree Star, Ashland, OR). The signal obtained with 293T (GFP-population) was subtracted from the signal obtained with 293T-Spike (GFP+ population) to remove unspecific signal.

### Neutralization assay using pseudoviral particles

Target cells were infected with single-round luciferase-expressing lentiviral particles as described previously (Prévost et al., 2020). Briefly, 293T cells were transfected by the calcium phosphate method with the lentiviral vector pNL4.3 R-E-Luc (NIH AIDS Reagent Program) and a plasmid encoding for SARS-CoV-2 Spike at a ratio of 5:4. Two days post-transfection, cell supernatants were harvested and stored at −80°C until use. 293T-ACE2 target cells were seeded at a density of 1×10^4^ cells/well in 96-well luminometer-compatible tissue culture plates (Perkin Elmer) 24h before infection. Recombinant viruses in a final volume of 100μl were incubated with the indicated plasma dilutions (1/50; 1/250; 1/1250; 1/6250; 1/31 250) for 1h at 37°C and were then added to the target cells followed by incubation for 48h at 37°C; cells were lysed by the addition of 30μl of passive lysis buffer (Promega) followed by one freeze-thaw cycle. An LB941 TriStar luminometer (Berthold Technologies) was used to measure the luciferase activity of each well after the addition of 100μl of luciferin buffer (15mM MgSO_4_, 15mM KPO_4_ [pH 7.8], 1mM ATP, and 1mM dithiothreitol) and 50μl of 1mM d-luciferin potassium salt (Prolume). The neutralization half-maximal inhibitory dilution (ID_50_) represents the sera dilution to inhibit 50% of the infection of 293T-ACE2 cells by recombinant viruses.

### Microneutralization assay using live SARS-CoV-2 viral particles

A microneutralization assay for SARS-CoV-2 serology was performed as previously described (Amanat et al., 2020). The assay was conducted with the person blinded to the sample identity. Experiments were conducted with the SARS-CoV-2 USA-WA1/2020 virus strain. This reagent was deposited by the Centers for Disease Control and Prevention and obtained through BEI Resources, NIAID, NIH: SARS-Related Coronavirus 2, Isolate USA-WA1/2020, NR-52281. One day prior to infection, 2×10^4^ Vero E6 cells were seeded per well of a 96 well flat bottom plate and incubated overnight (37°C/5% CO_2_) to permit Vero E6 cell adherence. On the day of infection, all plasma samples were heat inactivated at 56°C for one hour. Non-depleted plasma from each donor was also included in this assay. Plasma dilutions were performed in a separate 96 well culture plate using MEM supplemented with penicillin (100 U/mL), streptomycin (100 μg/mL), HEPES, L-Glutamine (0.3 mg/mL), 0.12% sodium bicarbonate, 2% FBS (all from Thermo Fisher Scientific) and 0.24% BSA (EMD Millipore Corporation). Plasma dilutions ranged from 1:50 to 1:31 250. In a Biosafety Level 3 laboratory (ImPaKT Facility, Western University), 10^3^ TCID_50_/mL SARS-CoV-2 USA-WA1/2020 virus strain was prepared in MEM + 2% FBS and combined with an equivalent volume of respective plasma dilution for one hour at room temperature. After this incubation, all media was removed from the 96 well plate seeded with Vero E6 cells and virus:plasma mixtures were added to each respective well at a volume corresponding to 600 TCID_50_ per well and incubated for one hour further at 37°C. Both virus only and media only (MEM + 2% FBS) conditions were included in this assay. All virus:plasma supernatants were removed from wells without disrupting the Vero E6 monolayer. Each plasma dilution (100 μL) was added to its respective Vero E6-seeded well in addition to an equivalent volume of MEM + 2% FBS and was then incubated for 48 hours. Media was then discarded and replaced with 10% formaldehyde for 24 hours to cross-link Vero E6 monolayer. Formaldehyde was removed from wells and subsequently washed with PBS. Cell monolayers were permeabilized for 15 minutes at room temperature with PBS + 0.1% Triton X-100 (BDH Laboratory Reagents), washed with PBS and then incubated for one hour at room temperature with PBS + 3% non-fat milk. An anti-mouse SARS-CoV-2 nucleocapsid protein (Clone 1C7, Bioss Antibodies) primary antibody solution was prepared at 1 μg/mL in PBS + 1% non-fat milk and added to all wells for one hour at room temperature. Following extensive washing with PBS, an anti-mouse IgG HRP secondary antibody solution was formulated in PBS + 1% non-fat milk. One hour post-room temperature incubation, wells were washed with PBS, SIGMA*FAST*™ OPD developing solution (Millipore Sigma) was prepared as per manufacturer’s instructions and added to each well for 12 minutes. Dilute HCl (3.0 M) was added to quench the reaction and the optical density at 490 nm of the culture plates was immediately measured using a Synergy LX multi-mode reader and Gen5™ microplate reader and imager software (BioTek®).

### Statistical analysis

Statistics were analyzed using GraphPad Prism version 8.0.2 (GraphPad, San Diego, CA, (USA). Every data set was tested for statistical normality and this information was used to apply the appropriate (parametric or nonparametric) statistical test. P values <0.05 were considered significant; significance values are indicated as *p<0.05; **p<0.01; ***p<0.001; ****p<0.0001.

